# *In Vitro* Studies of Antioxidant, Antidiabetic, and Antibacterial Activities of *Leea rubra* Blume

**DOI:** 10.1101/2024.11.26.625497

**Authors:** Van Mai Do, Truong Han Le, Hong Phong Ngo, Phat Nguyen Phung, Van Hung Mai

## Abstract

The current work investigated the antioxidant, antibacterial, and antidiabetic properties of extracts from *Leea rubra* roots, stems, and leaves *in vitro*. The antioxidant capabilities of *Leea rubra* extracts were tested using 2,2-diphenyl-1-picryl-hydrazyl, 2,2′-azino-bis (3-ethylbenzthiazoline-6-sulfonic acid), nitric oxide, total antioxidant activity, ferric reducing-antioxidant power, and reducing power tests. The extracts′ antibacterial activity against *Pseudomonas aeruginosa, Staphylococcus aureus*, and *Escherichia coli* was tested using the agar-well diffusion method. The antidiabetic efficacy of the extract was assessed by inhibiting α-amylase and α-glucosidase. *Leea rubra* extracts have shown strong antioxidant and antidiabetic action, with IC_50_ values ranging from 6.63±0.05 to 27.49±0.64 μg/mL and 20.31±0.46 to 41.84±0.02 μg/mL, respectively. *Leea rubra* extracts demonstrated significant antibacterial activity, with minimum inhibitory concentrations ranging from 160 to 640 μg/mL. Furthermore, the chemical composition study revealed the existence of alkaloids, polyphenols, flavonoids, steroids, tannins, saponins, and glycosides. This study reveals that *Leea rubra* contains a high potential of natural antibacterial, antioxidant, and antidiabetic properties.

## INTRODUCTION

Excessive production of free radicals, such as reactive oxygen species or reactive nitrogen species, from the environment and reactions/ metabolism in the human body leads to chronic diseases. Antioxidants play an important role in controlling the activity of free radicals (Jomova et al., 2023). Long-term usage of synthetic antioxidants can pose significant health hazards, hence they are frequently replaced by natural antioxidants in diets and health supplements. Uncontrolled free radical activity is strongly associated with diabetes. As a result, evaluating antioxidant activity is critical for identifying prospective antidiabetic medication sources. The increase of glucose in the blood causes oxidized glucose to reactive oxygen species during the glycation process, causing damage to different tissues and organs such as the eyes, kidneys, and heart, as well as increasing the level of lipid peroxidation in the body (González et al. 2023). Carbohydrate-metabolizing enzymes, including α-amylase and α-glucosidase, break down sugar and raise blood sugar levels. Plant secondary metabolites with antioxidant, α-amylase, and α-glycosidase inhibitory properties may help manage diabetes.

Multidrug-resistant bacteria pose a global health and economic risk. The emergence of multidrug-resistant bacteria has greatly lowered the efficacy of antimicrobial weaponry and raised the likelihood of treatment failure (Bharadwaj et al., 2022) (Muteeb et al., 2023). The problem of multidrug resistance in bacteria to synthetic antibiotics has shifted scientists′ focus to the discovery of novel natural antibacterial substances. Secondary metabolites from regularly used plants could be a source of novel antimicrobials (Murugaiyan et al., 2022). People in Vietnam have a long history of using plants as medicine to treat infectious infections, inflammatory conditions, injuries, and other maladies.

Plants of the genus *Leea* are traditionally used to treat various ailments such as fever, diarrhea, dysentery, arthralgia, rheumatism, diabetes, bone fractures, body aches, wounds, and sexual disorders. Most species of the genus *Leea* are medicinal plants with anti-cancer, cytotoxic, antibacterial, anti-diabetic, hepatoprotective, and cardiovascular activities (Hossain et al., 2021). *Leea rubra* is a plant in Vietnam that belongs to the genus *Leea*, which can grow in a variety of soils, is drought tolerant, and has formed roots that swell into tubers. *Leea rubra* leaves have been shown to contain important biological activities such as antibacterial, antitumor, antioxidant, and anti-inflammatory properties (Das et al., 2021). *Leea rubra* leaf extract contains numerous secondary metabolites, including polyphenols and flavonoids (Das et al., 2022). To our knowledge, our study is the most comprehensive study comparing the bioactivities among extracts from different parts of *Leea rubra*. The results of this study can be used to verify the pharmacological effects and can be used as a source of potential new antibacterial, antioxidant, and antidiabetic drugs.

## MATERIALS AND METHODS

### Research object

*Leea rubra* was collected in April 2024 in Long Trung commune, Cai Lay district, Tien Giang province, Vietnam (coordinates 10°19′34.2”N 106°06′08.9”E). *Leea rubra* was identified based on morphological characteristics as described in the book Dictionary of Vietnamese Medicinal Plants by Vo Van Chi (2021) with the support of Dr. Thieu Van Duong (Head of Biochemistry Department, Tay Do University). The morphological characteristics of *Leea rubra* are presented in Figure 1.

**Figure 1.**
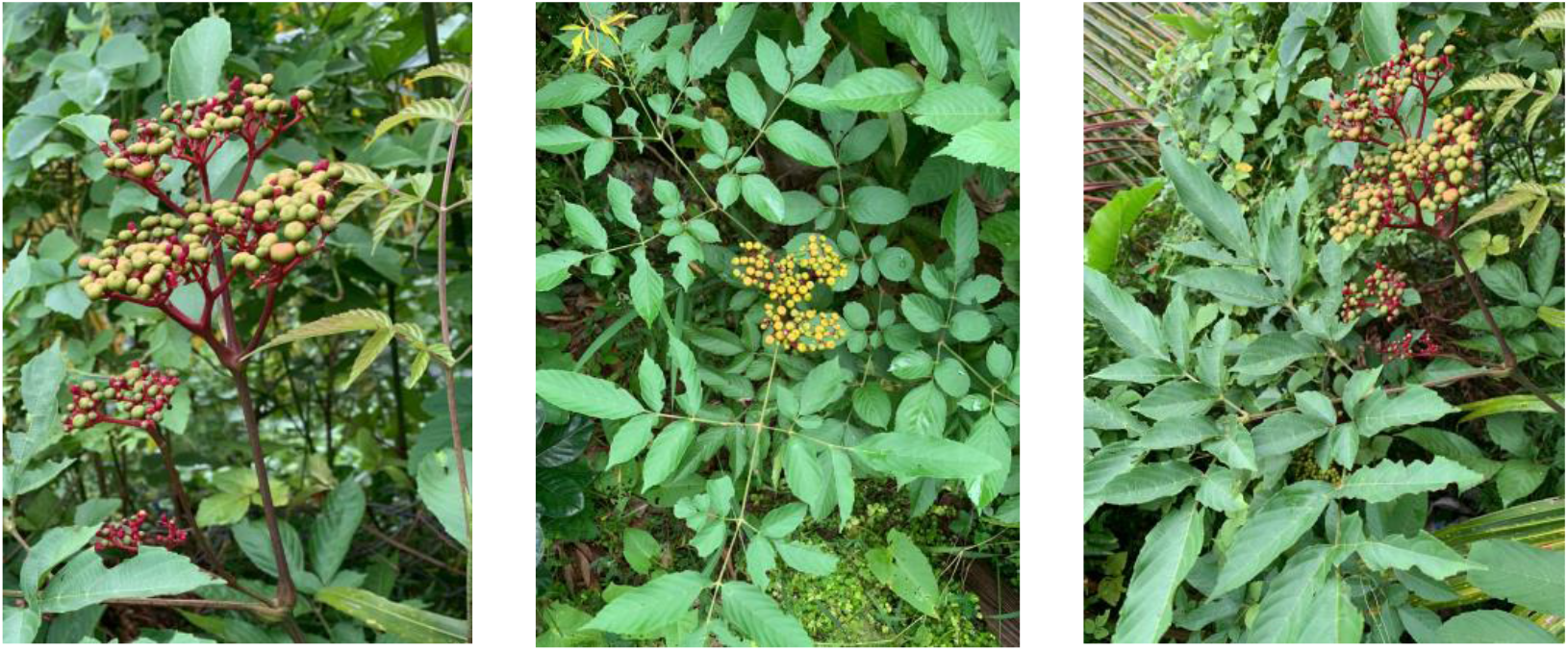
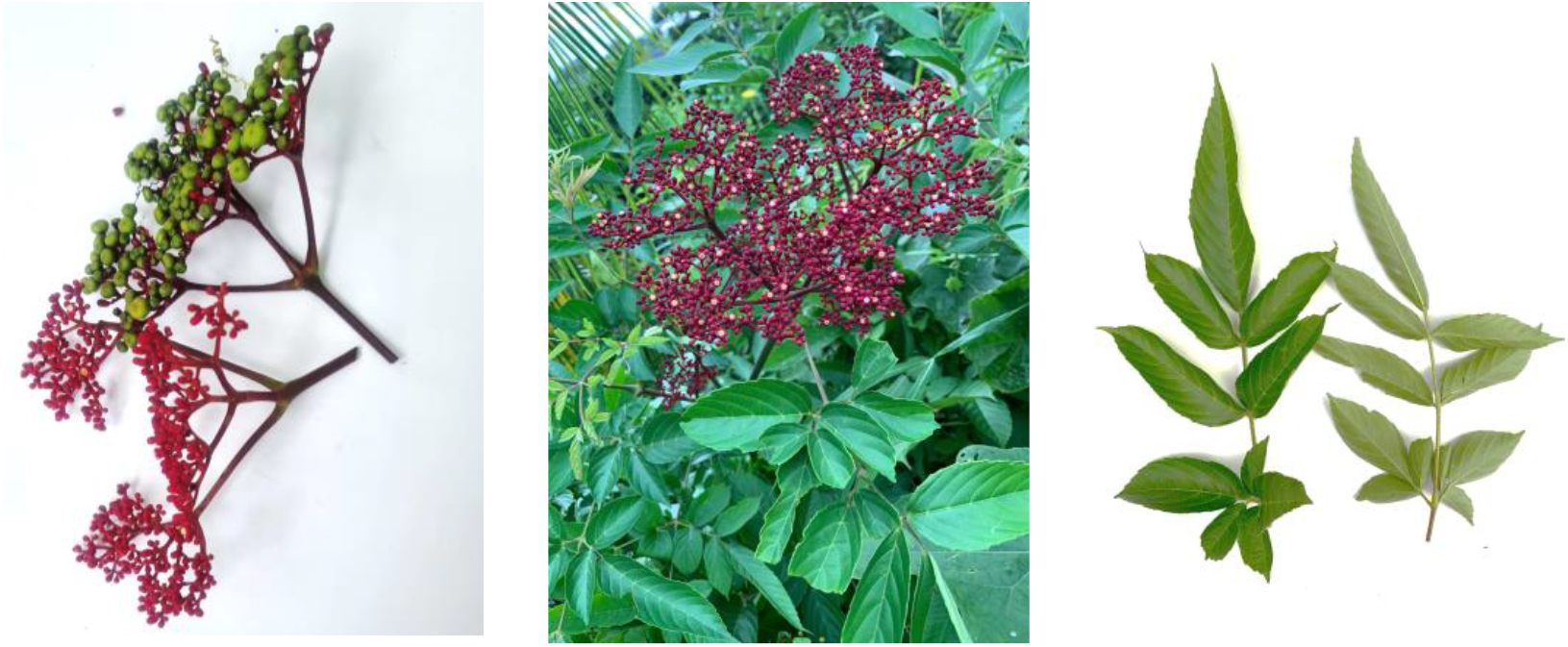
The morphology of *Leea rubra*

Equipment: Drying oven (BE 200, Memmert, Germany), analytical balance (AB104-S, Mettler Toledo, Switzerland), cold centrifuge (Mikro 12-24, Hettich, Germany), incubation tank (Memmert, Germany), rotary vacuum evaporator (Heidolph, Germany), spectrophotometer (Thermo Scientific Multiskan GO, Finland), autoclave sterilizer Sturdy SA-300VF (STURDY, Taiwan), biological safety cabinet Jeiotech BC-11B (Jeiotech, Korea), and advanced vortex mixer ZX3 (Velp, Italia).

Chemicals: Ethanol is offered by Cemaco (Vietnam). Folin-Ciocalteu′s phenol reagent, sodium carbonate, dimethyl sulfoxide, potassium persulphate, potassium ferricyanide, methanol, trichloroacetic acid, gallic acid quercetin are offered by Merck (Germany). Ascorbic acid, 2,2-azino-bis (3-ethylbanzthiazoline-6-sulphonic acid), 2, 4, 6-tripyridyl-s-triazine, 2,2-diphenyl-1-picrylhydrazyl, ferric chloride, α-amylase enzyme, α-glucosidase enzyme, acarbose are provided by Sigma-Aldrich (United States). Sodium nitrite, aluminum chloride hexahydrate, ammonium heptamolybdate tetrahydrate are offered by Xilong (China).

### Method of preparing ethanol extract from *Leea rubra*

After collection, *Leea rubra* is cleaned and split into three parts: root, stem, and leaf. Then, each portion will be dried at a temperature of around 50°C and processed into a medicinal powder with a particle size of about 60 mesh. The moisture content of *Leea rubra* medicinal powders made from its roots, stems, and leaves is determined by employing heat to evaporate all of the water. The medicinal powder will be steeped in 96% ethanol at a 1/10 (w/v) ratio at room temperature for 24 hours. The medicinal powder is soaked three times, the extracts from each soaking period are collected, and the solvent is evaporated using a vacuum rotary evaporator under reduced pressure at 50°C to get concentrated extracts. Extracts from *Leea rubra* roots, stems, and leaves are preserved in glass bottles, labeled, and refrigerated at Nam Can Tho University′s Faculty of Pharmacy in Vietnam.

### Qualitative analysis of the chemical compounds in extracts

As Biswas et al. (2012) describe, the research team investigated qualitative compound classes including polyphenols, alkaloids, flavonoids, steroids, glycosides, saponins and tannins.

### Polyphenol and flavonoid content measurements in extracts

With a few modifications, the total polyphenol content was calculated in accordance with Patra et al. (2019). Sodium carbonate solution (500 μL, 10%) and 500 μL of the Folin-Ciocalteu reagent (20%) were used to treat the extracts. After that, the reaction mixture was left to incubate for 30 minutes at 40°C in the dark. Lastly, the reaction mixture′s spectral absorbance at room temperature at 765 nm was determined. Using the standard curve equation y = 0.0152x -0.051 (R^2^=0.9896), the total polyphenol concentration is reported as mg gallic acid (GAE) per 1 g extract.

With a few modifications, the methodology described by Akanni et al. (2014) was used to determine the total flavonoid content. 500 μL of the extract were kept at room temperature for five minutes before reacting with 100 μL sodium nitrite (5%). Then, 100 μL aluminum chloride hexahydrate (10%) is added and thoroughly shaken. Following a 5-minute incubation period at room temperature, the reaction mixture was supplemented with 1000 μL of 1 M sodium hydroxide and 800 μL deionized water. The reaction mixture′s spectral absorbance at 510 nm was lastly measured. Quercetin (QE) concentration per 1 g extract is expressed as mg of flavonoids based on the standard curve equation (y = 0.0074x + 0.0037 (R^2^ = 0.9998).

### Antioxidant efficacy of extracts *in vitro*

With some adjustments, the DPPH free radical neutralization method described by Xie et al. (2014) was used to assess the 2,2-diphenyl-1-picrylhydrazyl (DPPH) free radical neutralization activity of *Leea rubra* extracts. The reaction mixture contained 40 μL of DPPH (1000 μg/mL) and 960 μL of extract. This mixture was incubated at 30°C in the dark for thirty minutes. Then, the spectral absorbance of DPPH was measured at wavelength 517 nm.

Using the ABTS decolorization method developed by Ilyasov et al. (2020), the free radical neutralizing activity of 2,2′-azino-bis (3-ethylbenzthiazoline-6-sulfonic acid) (ABTS) was determined. A solution of 2.45 mM potassium persulfate and 7 mM ABTS was prepared. Before being used, the mixture was incubated for 16 hours at room temperature in the dark. Upon diluting the mixture, 0.70 ± 0.05 was found to be the spectral absorbance at 734 nm. Then, 10 μL of the extract was combined with 990 μL of ABTS at room temperature for 6 minutes to carry out the survey. Next, a measurement of the reaction mixture′s spectral absorbance at 734 nm was made.

With modifications, Griffin & Bhagooli′s (2004) description was used to calculate the reduction potential of *Leea rubra* extracts. This method′s reduction of the ferric-tripyridyltriazine complex serves as its foundation. After mixing 10 μL of *Leea rubra* extract with 990 μL of FRAP solution for 30 minutes in low light, the mixture was left to settle. The spectral absorbance of the experimental solution was measured at 593 nm.

The total antioxidant activity of *Leea rubra* extracts were assessed using the method published by Umamaheswari et al. (2007). The *Leea rubra* extract (300 μL) was mixed with 900 μL test solution (0.6 M sulfuric acid, 28 mM sodium phosphate, and 4 mM ammonium molybdate). For 90 minutes, the reaction solution was incubated at 95°C. The solution′s spectral absorbance was measured at 695 nm.

According to Alisi & Onyeze (2007), with modifications, the capacity of the *Leea rubra* extract to suppress the generation of nitric oxide (NO) was studied. The extract (200 μL) and 400 μL of sodium nitroprusside (5 mM) were added to the reaction mixture. The reaction mixture was centrifuged for 15 minutes at 11.000 rpm after being incubated for 60 minutes at 25°C. There was an additional 600 μL of Griess reagent added to the centrifuge. After an additional five minutes of incubation, the sample was examined for spectral absorbance at 546 nm.

In the methods for assessing antioxidant activity outlined above, essential ascorbic acid functioned as a positive control. The extract from *Leea rubra* was tested in vitro for antioxidant activity against a standard of ascorbic acid, using a concentration (μg/mL) that reduced, neutralized, or blocked 50% of free radicals (IC50-inhibitory concentration of 50%). The IC50 values for ascorbic acid and *Leea rubra* extract were determined following the findings of Piaru et al. (2008).

### Examination of extracts′ *in vitro* antidiabetic potential

With few adjustments, the α-amylase inhibitory activity of *Leea rubra* extracts was assessed as previously reported by Mohamed et al. (2012). The reaction mixture was made up of 100 μL of extract at various concentrations and 100 μL of pH 7 phosphate buffer that was treated for five minutes at 37 °C with 100 μL of starch (2 mg/mL). After being added to the reaction mixture, the α-amylase enzyme (3 U/mL, 100 μL) was incubated for 15 minutes at 37 °C. A final step was adding 400 μL of 1 M hydrochloric acid to halt the process. After adding 600 μL of iodine reagent, the reaction mixture′s spectral absorbance was measured at 660 nm.

With certain changes, the α-glucosidase enzyme inhibitory activity of *Leea rubra* extracts was found, as reported by Chipiti et al. (2015). The extracts (250 μL) of *Leea rubra* were subjected to a 15-minute incubation period at 37°C with 500 μL of α-glucosidase enzyme (1 U/mL) mixed in 100 mM phosphate buffer (pH 6.8). The mixture was then incubated at 37°C for 20 minutes after 250 μL of 4-nitrophenyl-D-glucopyranoside solution (5 mM, diluted in 100 mM phosphate buffer; pH 6.8) was added. At room temperature, the spectral absorbance of the p-nitrophenol produced during the reaction was measured at 405 nm.

As stated by Mohamed et al. (2012), the capacity to suppress α-amylase and α-glucosidase enzyme activity was assessed using inhibition effectiveness (%) and concentration (μg/mL) that suppresses 50% of enzyme activity (often referred to as the IC_50_ value-inhibitory concentration of 50%). Furthermore, acarbose was compared to the inhibitory effects of *Leea rubra* extracts against α-amylase and α-glucosidase.

### Antibacterial properties of extracts *in vitro*

The study used *Escherichia coli* NCTC 13216, *Pseudomonas aeruginosa* ATCC^®^ 27853™, and *Staphylococcus aureus* ATCC^®^ 25923™ bacteria. These bacterial strains were cultured and stored at the Faculty of Medicine of Nam Can Tho University. The antibacterial activity was tested using the agar well diffusion technique, as reported by Ngan et al. (2012). 0.01 g of each extract was dissolved in 10% DMSO (1,000 μL) to form the concentration of 10,000 μg/mL. The extracts were then diluted using 10% DMSO solvent to solutions with concentrations of 80, 160, 320, 640, and 1280 μg/mL. The bacterial solution showed an optical density of 0.5 (OD_600_ = 0.5) at a wavelength of 600 nm after being diluted in physiological saline. After applying 100 μL of the bacterial solution to the surface and drying it, five 7 mm-diameter wells were punched on a petri dish filled with Luria Bertani (LB) agar media. To test the bacterial suspension, 50 μL of extract at different concentrations was added to the wells of a petri dish filled with LB agar medium. The test sample is incubated for twenty-four hours at 37°C. Tthe diameter of the antibacterial zone that forms around the wells on the agar plate was measured and recorded after a full day had passed.

### Data analysis statistics

Statistical analysis: Using the minitab 16 for Windows tool, one-way ANOVA (Tukey′s studentized range) was performed on the mean data, which were reported as Mean±Standard Deviation (SD). When p < 0.05, differences were deemed significant.

## RESULTS

### Results of preparation, qualitative and quantitative determination of chemical components in the extract

*Leea rubra*′s fresh roots (900 g), stems (900 g), and leaves (900 g) were harvested, dried, and crushed into powdered medicinal herbs: roots (185 g), stems (190 g), and leaves (150 g), with moisture levels of 10.6%, 11.4%, and 10.2%, respectively. Through the process of preparing the extract, we obtained the roots, stems, and leaves extracts of *Leea rubra* with weights of 8.5 g, 9.6 g, and 9.0 g, respectively. The qualitative chemical composition of *Leea rubra* roots, leaves, and stems revealed the presence of alkaloids, polyphenols, flavonoids, and glycosides. Additionally, the root and stem preparations of *Leea rubra* contained steroids. The leaf extract of *Leea rubra* included tannin and saponin components. Table 1 shows the qualitative chemical makeup of *Leea rubra* extracts.

**Table 1.** The qualitative results of the chemical composition of the extract.

| Phytochemicals | Test | Inference |  |  |
| --- | --- | --- | --- | --- |
|  |  | Root | Leaf | Stem |
| Alkaloids | Dragendroff | + | + | + |
|  | Wagner | + | + | + |
| Polyphenols | Folin-Ciocalteu | + | + | + |
| Flavonoids | FeCl<sub>3</sub> 5% | + | + | + |
|  | 1% NaOH/ethanol | + | + | + |
| Triterpenoids | Rosenthaler | — | — | — |
| Steroids | Salkowski | + | — | + |
| Tannins | Pb(CH<sub>3</sub>COO)<sub>2</sub><br>(Saturated) | — | + | — |
| Saponins | NaOH 0,1N (pH = 13) | — | — | — |
|  | HCl 0,1N (pH = 1) | — | + | — |
| Glycosides | Fehling | + | + | + |

Note: (+) denotes presence and (-) denotes absence in the extracts.

Figure 2 shows the polyphenol and flavonoid content of the three *Leea rubra* extracts. The polyphenol content of the various extracts ranged between 153.40±5.81 and 188.27±11.94 mg GAE/g extract. The root extract has the highest total flavonoid concentration (105.17±4.22 mg QE/g extract), followed by the stem extract (92.22±0.52 mg QE/g extract) and the leaf extract (79.49±1.88 mg QE/g extract).

**Figure 2.**
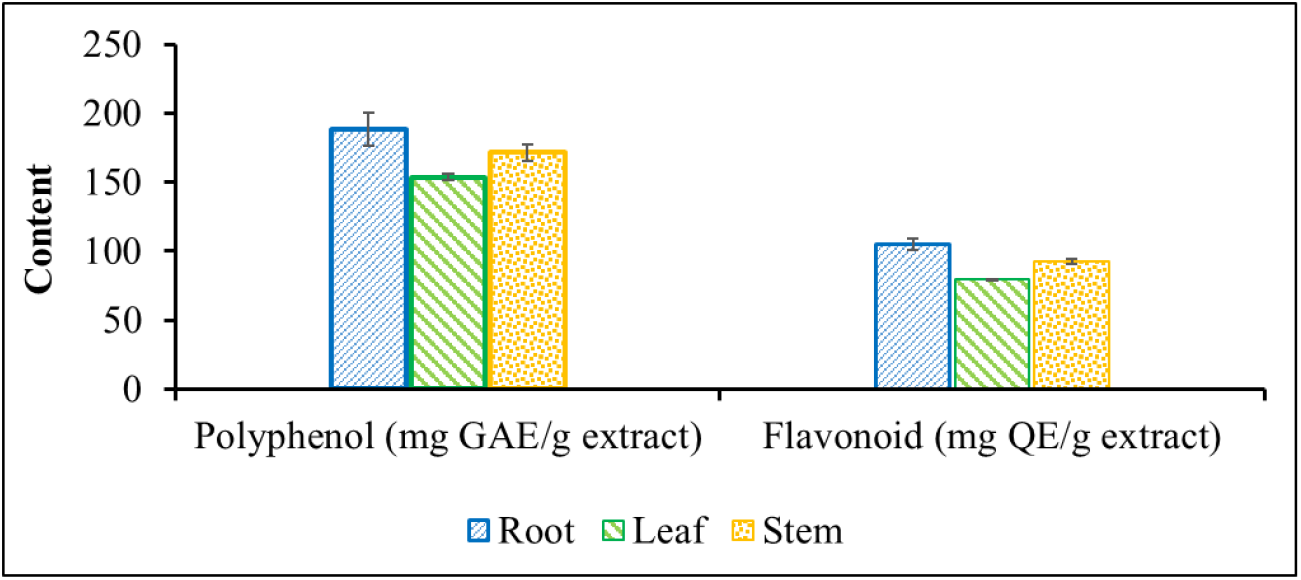
Total polyphenol and flavonoid contents of *Leea rubra* extracts

### The antioxidant properties of extracts

The free radical neutralization efficiency of ABTS^·+^, DPPH, and NO^·^ is shown in Figure 3. All The extracts from *Leea rubra* have the ability to neutralize free radicals ABTS, DPPH, and NO. Among that, the roots extract illustrated the highest efficiency (ABTS^·+^: 5.40±0.60–66.64±0.62%, DPPH: 6.23±0.52–77.19±0.72%, NO^·^: 14.11±1.56–86.17±1.07%), followed by the stems (ABTS^·+^: 6.46±1.05–57.38±1.18%, DPPH: 11.63±0.24– 67.49±2.07%, NO^·^: 7.79±1.66–66.26±1.11%), and finally the leaves (ABTS^·+^: 5.44±0.52–53.24±0.95%, DPPH: 7.47±0.69–53.51±1.22%, NO^·^: 7.45±1.68–55.31±1.73%). The free radical neutralization efficiency of ABTS^·+^, DPPH, and NO^·^ of the extracts from *Leea rubra* increased gradually with the concentration of the test sample. In this study, the IC_50_ values for ABTS^·+^, DPPH, and NO^·^ of *Leea rubra* root extract were 7.66±0.19, 6.63±0.05, and 18.08±0.41 μg/mL, respectively, indicating the highest free radical scavenging activity. In contrast, *Leea rubra* leaf extract showed the lowest free radical scavenging activity with IC_50_ values for ABTS, DPPH, and NO of 9.29±0.08, 8.98±0.21, and 27.49±0.64 μg/mL, respectively. The ABTS^·+^, DPPH, and NO^·^ free radical scavenging activity of *Leea rubra* root extract was 1.21, 1.35, and 1.52 times stronger than that of *Leea rubra* leaf extract, respectively. *Leea rubra* root and stem extracts had 1.11 and 1.05 times stronger DPPH free radical scavenging activity than ascorbic acid, respectively. Ascorbic acid had 3.79, 2.97, and 2.50 times weaker NO^·^ free radical scavenging activity than *Leea rubra* root, stem, and leaf extracts, respectively.

**Figure 3.**
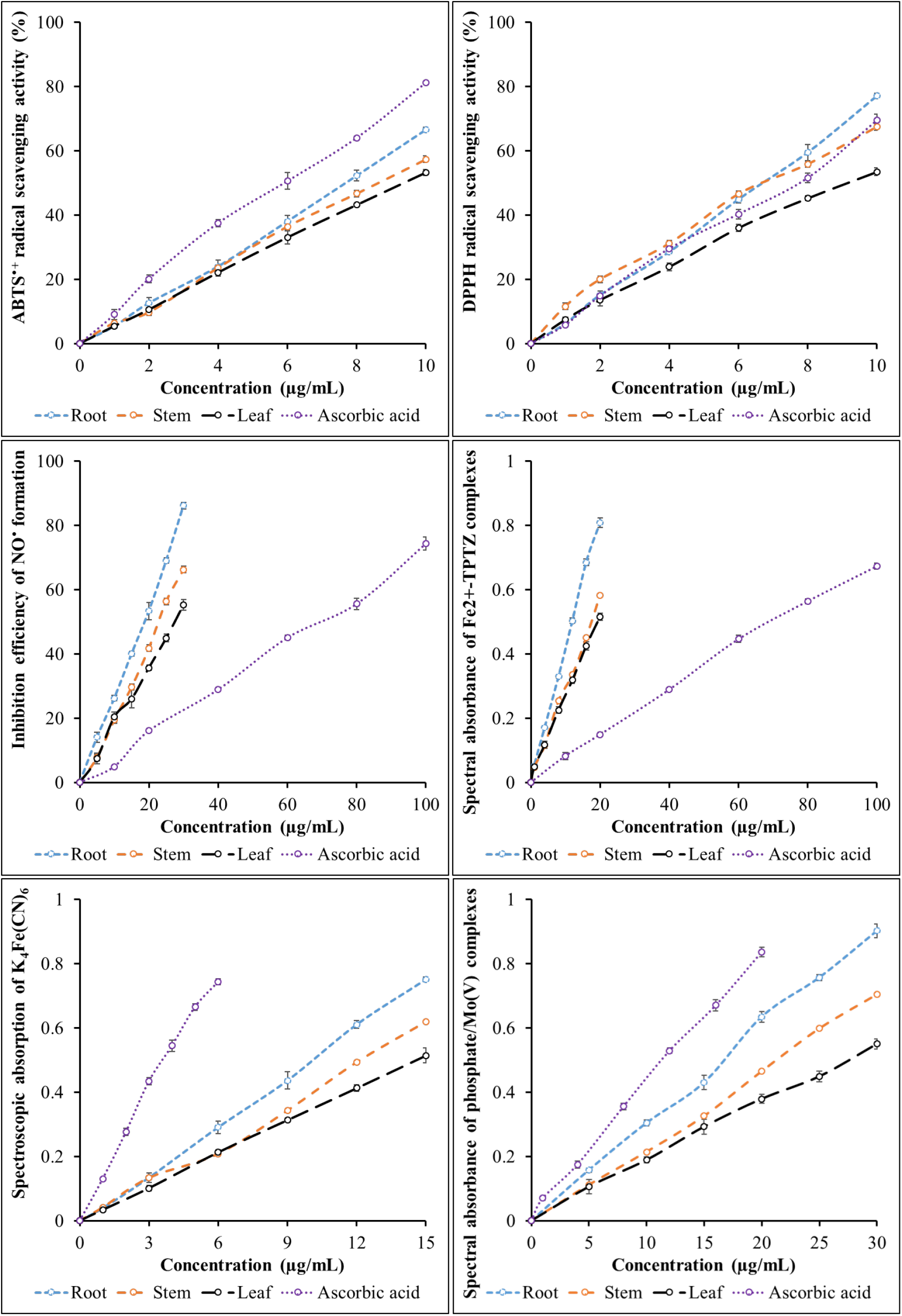
Neutralization/inhibition efficiency or spectral absorption of free radicals

The total antioxidant activity of *Leea rubra* extracts was determined based on the reduction of Mo(VI) to Mo(V) by secondary metabolites. The spectral absorbance of the phosphate/Mo(V) complex was determined and presented in Figure 3F. The spectral absorbance of the phosphate/Mo(V) complex increased gradually with the concentration of the test sample (root: 0.158±0.007-0.902±0.022; stem: 0.112±0.022-0.705±0.009; and leaf: 0.106±0.023-0.550±0.016). Among them, the root extract of *Leea rubra* had the strongest total antioxidant activity. The root (IC_50_=16.47±0.25 μg/mL), stem (IC_50_=21.44±0.25 μg/mL), and leaf (IC_50_=21.44±0.25 μg/mL) extracts of *Leea rubra* had 1.41, 1.83, and 2.32 times weaker total antioxidant activity than ascorbic acid (IC_50_=11.69±0.09 μg/mL), respectively.

In RP and FRAP methods, Fe^3+^ complex was reduced to Fe^2+^ complex by secondary metabolites present in *Leea rubra* roots, stems, and leaves. Fe^3+^ reduction activity increased gradually with the concentration of the extracts (Figures 3D, 3E). *Leea rubra* extracts were all able to reduce Fe^3+^ complex to Fe^2+^ complex with the highest spectral absorption in roots (RP: 0.039±0.007–0.752±0.008, FRAP: 0.053±0.005–0.808±0.015), followed by stems (RP: 0.042±0.012–0.620±0.015, FRAP: 0.048±0.005–0.583±0.015) and finally leaves (RP: 0.034±0.006– 0.515±0.023, FRAP: 0.048±0.004–0.516±0.011). The IC_50_ values of the extracts based on RP and FRAP assays are shown in Table 2. In RP and FRAP methods, the antioxidant activity increased in the order from leaves (IC_50, RP_=14.48±0.21 μg/mL; IC_50, FRAP_=19.12±0.41 μg/mL) then the stems (IC_50, RP_=12.38±0.14 μg/mL; IC_50, FRAP_=17.39±0.34 μg/mL), and the roots is the most effective (IC_50, RP_=10.04±0.20 μg/mL; IC_50, FRAP_=12.02±0.16 μg/mL).

**Table 2.** Antioxidant activity of extracts.

| Sample | The IC<sub>50</sub> value (µg/mL) |  |  |  |  |  |
| --- | --- | --- | --- | --- | --- | --- |
|  | ABTS<sup>•+</sup> | DPPH | NO<sup>•</sup> | TAC | RP | FRAP |
| Root | 7.66±0.19 | 6.63±0.05 | 18.08±0.41 | 16.47±0.25 | 10.04±0.20 | 12.02±0.16 |
| Stem | 8.59±0.26 | 6.99±0.16 | 23.08±0.47 | 21.44±0.25 | 12.38±0.14 | 17.39±0.34 |
| Leaf | 9.29±0.08 | 8.98±0.21 | 27.49±0.64 | 27.17±0.93 | 14.48±0.21 | 19.12±0.41 |
| Ascorbic acid | 6.00±0.06 | 7.35±0.11 | 68.59±0.60 | 11.69±0.09 | 3.79±0.05 | 71.49±0.11 |

Note: Different letters in the same column show significant difference at level of 5%.

**Table 3.** The α-amylase and α-glucosidase inhibitory activities of the extracts.

| Sample | Giá trị $IC_{50}$ ($\mu\text{g/mL}$) | |
| --- | --- | --- |
| | α-Amylase | α-Glucosidase |
| Root | $35.03\pm0.42$ | $20.31\pm0.46$ |
| Stem | $38.15\pm0.81$ | $23.76\pm0.39$ |
| Leaf | $41.84\pm0.02$ | $25.20\pm0.23$ |
| Acarbose | $5.84\pm0.29$ | $3.55\pm0.02$ |

Note: Different letters in the same column show significant differences at level of 5%.

### *In vitro* antidiabetic activity of the extract

*Leea rubra* extracts showed *in vitro* antidiabetic efficacy by inhibiting α-amylase and α-glucosidase enzymes. Figure 4 illustrates how *Leea rubra* extracts block α-amylase and α-glucosidase enzymes. *Leea rubra* extracts inhibited α-amylase and α-glucosidase enzymes with efficiencies ranging from 3.97±0.98 to 82.00±2.29%. The inhibitory efficiency of α-amylase and α-glucosidase enzymes of *Leea rubra* extracts was also concentration dependent.

**Figure 4.**
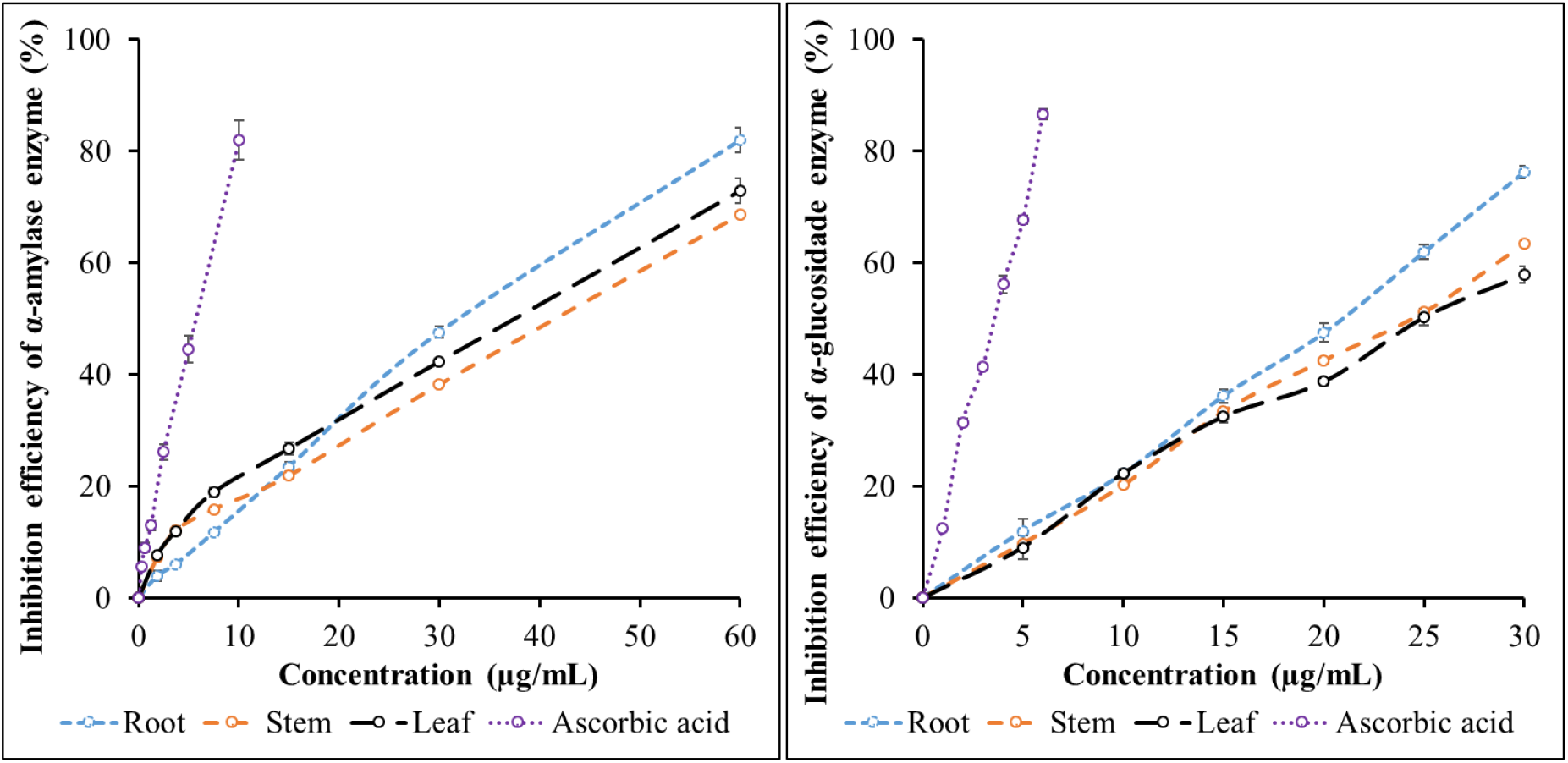
Inhibitory effects of α-amylase and α-glucosidase enzymes of the extracts

The strongest *in vitro* antidiabetic activity was determined in the roots (IC_50, α-amylase_=35.03±0.42 μg/mL; IC_50, α-glucosidase_=20.31±0.46 μg/mL), followed by the stems (IC_50, α-amylase_=38.15±0.81 μg/mL; IC_50, α-glucosidase_=23.76±0.39 μg/mL) and finally the leaves (IC_50, α-amylase_=41.84±0.02 μg/mL; IC_50, α-glucosidase_=25.20±0.23 μg/mL). Acarbose had IC50 values of 5.84±0.29 for α-amylase and 3.55±0.02 for α-glucosidase, respectively, indicating that acarbose is very effective in inhibiting α-amylase and α-glucosidase enzymes.

### Antibacterial activity of extracts

Figure 5 shows the diameter of the zone of inhibition, which demonstrates the antibacterial activity of *Leea rubra* extracts. *Leea rubra* extracts were found to inhibit *Escherichia coli, Pseudomonas aeruginosa*, and *Staphylococcus aureus*. The largest antibacterial zone diameters were found in root extracts (*Escherichia coli*: 15.03±1.10–21.27±0.65 mm, *Pseudomonas aeruginosa*: 14.67±1.11–21.13±0.29 mm, *Staphylococcus aureus*: 9.77±0.21–25.67±1.11 mm), followed by stems (*Escherichia coli*: 12.87±1.40–19.03±0.61 mm, *Pseudomonas aeruginosa*: 12.47±0.95–18.77±0.47 mm, *Staphylococcus aureus*: 7.90±0.76–23.47±1.10 mm), and finally leaves (*Escherichia coli*: 13.17±0.99–17.17±0.78 mm, *Pseudomonas aeruginosa*: 8.17±0.21–10.37±1.15 mm, *Staphylococcus aureus*: 7.90±0.79–21.57±0.32 mm).

**Figure 5.**
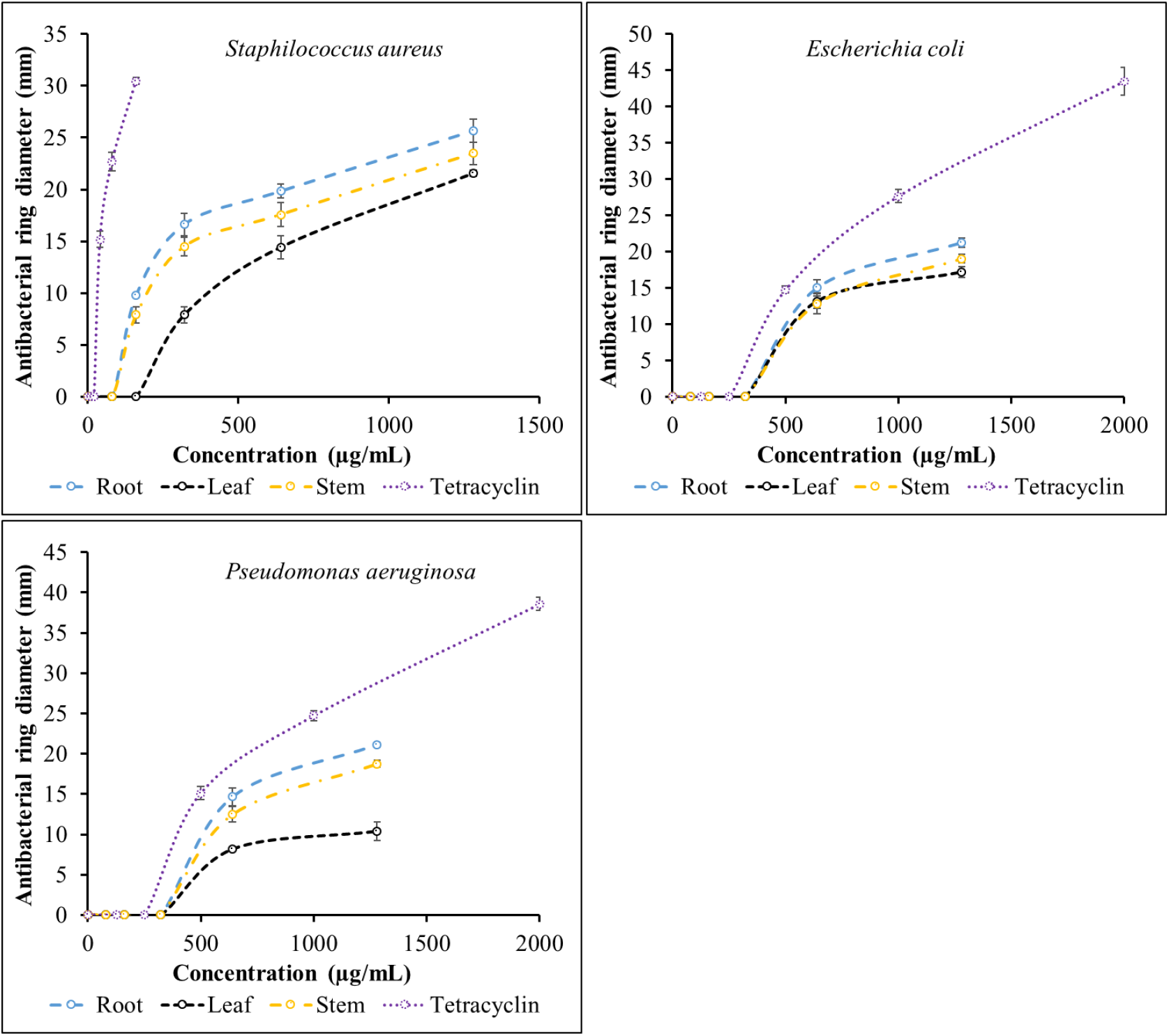
Antibacterial activity of extracts

The study also determined the minimum inhibitory concentrations of each extract against each bacterial strain (Table 4). The minimum inhibitory concentrations of *Leea rubra* extracts against *Escherichia coli, Pseudomonas aeruginosa*, and *Staphylococcus aureus* ranged from 160 to 640 μg/mL.

**Table 4.** Minimum inhibitory concentration.

| Sample | Giá trị MIC (µg/mL) |  |  |
| --- | --- | --- | --- |
|  | *Escherichia coli* | *Pseudomonas aeruginosa* | *Staphylococcus aureus* |
| Root | 640 | 640 | 160 |
| Stem | 640 | 640 | 160 |
| Leaf | 640 | 640 | 320 |
| Tetracyclin | 500 | 500 | 40 |

## DISCUSSION

In nature, to adapt to biotic and abiotic stress factors such as pathogens, high temperatures, salinity, and prolonged drought, plants must produce a variety of secondary metabolites (polyphenols, alkaloids, flavonoids, steroids, glysosides, saponins, and tannins). Secondary metabolites from plants provide important health benefits to humans. This has attracted the attention of scientists to investigate the presence of these secondary metabolites. In our study, extracts from *Leea rubra* possessed a variety of plant secondary metabolites with rich biological potential. However, we focused our attention on the polyphenol and flavonoid categories in our study. Because polyphenols and flavonoids are present in three extracts of *Leea rubra*, as well as the health benefits that polyphenols and flavonoids provide. Secondary metabolites from the polyphenol and flavonoid groups have antioxidant, anti-inflammatory, anticancer, antibacterial, body-protective, and disease-treating properties in many human disorders. The compound myricetin 4′-methoxy-3-O-α-l-rhamnopyranoside is a flavonoid extracted from *Leea rubra* leaves that has protective effects against DNA damage and anticancer effects (Das et al., 2022). In this study, the root extract of *Leea rubra* had a greater total polyphenol and flavonoid content than the stem and leaf extracts. Plant roots are more susceptible to environmental influences and infections than stems and leaves. This causes plant roots to create more secondary metabolites. This result is similar to the study of Uddin et al. (2022) on the content of polyphenols and flavonoids in roots > stems > leaves. In general, the total amount of polyphenols and flavonoids plays an important role in the antioxidant, α-amylase, α-glucosidase inhibitory, and antibacterial activities of the extract. Many studies have determined the dose-effect relationship between biological activity and total polyphenol and flavonoid content. However, for different parts of plants, there is not always a relationship between biological activities and total polyphenol and flavonoid content; it also depends on the composition of polyphenols and flavonoids, not just the measurement of the decisive content (Larit et al., 2019; Ullah et al., 2020).

Antioxidants derived from plants play a much more useful and effective role in combating free radical-related diseases. Compared to synthetic chemicals, products derived from natural sources are safer for long-term use. Among the groups of natural compounds with antioxidant properties, polyphenols and flavonoids show many advantages. The antioxidant activity of compounds belonging to the polyphenol and flavonoid groups depends on their content and molecular (or chemical) structure. In this study, six antioxidant methods, including ABTS^·+^, DPPH, NO^·^, TAC, FRAP, and RP, were used to evaluate the antioxidant activity of *Leea rubra* extracts. ABTS^·+^ and DPPH are abiotic free radicals that are often selected in screening tests for new drugs with antioxidant activity. In FRAP and RP methods, the reducing power of secondary metabolites present in the extract is measured by reducing Fe^3+^ ions to Fe^2+^ ions. Electron donation potential is usually measured by iron reduction, and these are also suitable techniques to evaluate the antioxidant activity of polyphenols and flavonoids (Saleem et al., 2023). In the NO^·^ method, sodium nitroprusside in an aqueous solution at physiological pH produces NO^·^, which interacts with oxygen to form a nitrite ion, which is determined using Griess reagent. Total antioxidant activity is determined by the phosphomolybdenum assay based on the reduction of Mo(VI) to Mo(V), which is demonstrated by the formation of a blue phosphate/Mo(V) complex at acidic pH due to the antioxidant compounds present in the plant extract. This blue complex is quite stable for several days and is not affected by various organic solvents (Singh & Singh, 2008). In our study, the best antioxidant activity was detected in the roots of *Leea rubra*, followed in decreasing order by the stems and leaves of *Leea rubra* (Table 1). Polyphenols and flavonoids are polar substances and possess strong antioxidant activity that can neutralize free radicals by donating their hydrogen atoms and electrons. A positive correlation between the total polyphenol and flavonoid content in plant extracts and antioxidant activity was also observed in the studies of Muflihah et al. (2021) and Dibacto et al. (2021). The study by Joshi et al. (2016) showed that the root extract of *Leea macrophylla*, a plant species of the same genus as *Leea rubra*, also has the ability to neutralize DPPH free radicals with an IC_50_ value of 39.80±2.05 μg/mL. The roots, stems, and leaves of *Leea rubra* have the ability to neutralize DPPH free radicals stronger than the roots of *Leea macrophylla* by 6.00, 5.69, and 4.43 times, respectively. The extracts from *Leea rubra* have stronger antioxidant activity than the plants of the same genus.

*In vitro, Leea rubra* extracts inhibited carbohydrate metabolism by α-amylase and α-glucosidase enzymes, indicating potential antidiabetic action. The carbohydrate used in the study was starch. During metabolism, pancreatic α-amylase broke down starch into disaccharides and oligosaccharides. In the small intestine, α-glucosidase breaks down disaccharides into glucose (Oluwagunwa et al., 2021; Lam et al., 2024). Inhibiting the α-glucosidase enzyme slows starch digestion and glucose absorption in the digestive tract, leading to lower blood glucose levels. The *Leea rubra* extracts block α-amylase and α-glucosidase enzymes, with inhibition increasing from leaves to stems to roots (Table 1). The α-amylase and α-glucosidase inhibitory activities of *Leea rubra* extracts were similar to the investigated antioxidant activities. The ability to inhibit these enzymes correlated positively with the antioxidant capability of the extracts described above. Previous research indicated that plant-derived secondary metabolites with antioxidant action can also inhibit α-glucosidase and α-amylase. (Anh et al., 2021; Zakhele et al., 2023). Polyphenols have long been known for their inhibitory activity against α-amylase and α-glucosidase enzymes due to their specific molecular structure and content. Flavonoids are group of natural compounds belonging to the polyphenol group characterized by a C6-C3-C6 skeleton consisting of two benzene rings (rings A and B) linked via a three-carbon bridge. Notably, several studies have highlighted the potential of flavonoids as antidiabetic agents due to their strong inhibition of α-glucosidase and moderate inhibition of α-amylase. This is consistent with our study, extracts from *Leea rubra* were more effective in inhibiting α-glucosidase than α-amylase. According to Martinez-Gonzalez et al. (2019) and Barber et al. (2021), the ability of flavonoids to inhibit α-amylase and α-glucosidase enzymes depends on the position and number of hydroxyl groups in the molecule. Hydroxylation at the 3′ or 3 positions of flavone or at the 6, 3′, and 5′ positions of flavonols and isoflavones, as well as the 4′ position of flavanones, will enhance the inhibitory activity of flavonoids against enzymes because the hydroxyl group interacts with amino acid residues at the active sites of the enzyme.

The antibacterial activity of *Leea rubra* extracts was determined by the agar-well diffusion method. *Leea rubra* root extract had the strongest antibacterial activity, followed by stem extract, and finally *Leea rubra* leaf extract. This result is consistent with the polyphenol and flavonoid content analyzed above. *Leea rubra* extracts inhibited Gram-positive bacteria (*Staphilococcus aureus*) more effectively than Gram-negative bacteria (*Pseudomonas aeruginosa, Escherichia coli*). This may be related to the lipopolysaccharide membrane. Gram-positive bacteria are more sensitive due to the lack of an external lipopolysaccharide membrane. Gram-negative bacteria have an external lipopolysaccharide membrane and are less sensitive (Godoy-Gallardo et al., 2021). Many studies have shown that the antibacterial activity of antibacterial agents is related to many factors, including salt concentration, nutrient availability of pathogens, cell surface properties of bacteria, and the structure of plant secondary metabolites (Hochma et al., 2021; Zhang et al., 2022). Bacterial cell membranes play an important role in transport, protection, permeability, and cell biosynthesis, and cell membrane disruption can lead to bacterial death. Secondary metabolites of the polyphenol and flavonoid groups, due to the presence of hydroxyl groups, can form covalent bonds with transport proteins on the cell membrane, thereby increasing permeability and disrupting the cell membrane.

## CONCLUSION

The study determined the in vitro antioxidant, antibacterial, and antidiabetic activities of *Leea rubra* extracts. The results showed that *Leea rubra* root extract had stronger in vitro antioxidant, antibacterial, and antidiabetic activities than *Leea rubra* stem and leaf extracts. This was related to the polyphenol and flavonoid content in each extract. The polyphenol and flavonoid content in *Leea rubra* extracts increased gradually from leaves < stems < roots. *Leea rubra* extracts were more effective in inhibiting Gram-positive bacteria than Gram-negative bacteria. These *in vitro* biological activities contribute to explaining the uses of *Leea rubra* in folk medicine and the potential for exploiting active secondary metabolites from this plant.

## REFERENCES

Akanni, O.O., Owumi, S.E., Adaramoye, O.A., 2014. In vitro studies to assess the antioxidative, radical scavenging and arginase inhibitory potentials of extracts from Artocarpus altilis, Ficus exasperate and Kigelia africana. Asian Pacific Journal of Tropical Biomedicine, vol. 4, no. 1, pp. 492–499.

Alisi, C.S., Onyeze, G.O.C., 2008. Nitric oxide scavenging ability of ethyl acetate fraction of methanolic leaf extracts of Chromolaena odorata (Linn.). African Journal of Biochemistry Research, vol. 2, no. 7, pp. 145–150.

Anh, V.T.T., Trang, D.T.X., Kamei, K., Linh, T.C., Pham-Khanh, N.H., Tuan, N.T., Danh, L.T. Phytochemicals, antioxidant and antidiabetic activities of extracts from Miliusa velutina flowers, Horticulturae, vol. 7, p. 555.

Barber, E., Houghton, M.J., Williamson, G., 2021. Flavonoids as human intestinal α-glucosidase inhibitors. Foods, vol. 10, p. 1939.

Bharadwaj, A., Rastogi, A., Pandey, S., Gupta, S., Sohal, J.S., 2022. Multidrug-resistant bacteria: Their mechanism of action and prophylaxis. BioMed Research International, vol. 2022, p. 5419874.

Biswas, S.K., Chowdhury, A., Raihan, S.Z., Muhit, M.A., Akbar, M.A., Mowla, R., 2012. Phytochemical investigation with assessment of cytotoxicity and antibacterial activities of chloroform extract of the leaves of Kalanchoe pinnata. Journal of Plant Physiology, vol. 7, pp. 41–46.

Chipiti, T., Ibrahim, M.A., Singh, M., Islam, M.S., 2015. In vitro α-amylase and α-glucosidase inhibitory effects and cytotoxic activity of Albizia antunesiana extracts. Pharmacognosy Magazine, vol. 11, no. 2, pp. 231–236.

Das, N., Parvin, M.S., Hasan, M., Akter, M., Hossain, M.S., Parvez, G.M.M., Sarker, A.K., Abdur Rahman, M.A., Mamun, A., Islam, M.E., 2022. A flavone from the ethyl acetate extract of Leea rubra leaves with DNA damage protection and antineoplastic activity. Biochemistry and Biophysics Reports, vol. 30, p. 101244.

Dibacto, R.E.K., Tchuente, B.R.T., Nguedjo, M.W., Tientcheu, Y.M.T., Nyobe, E.C., Edoun, F.L.E., Kamini, M.F.G., Dibanda, R.F., Medoua, G.N., 2021. Total polyphenol and flavonoid content and antioxidant capacity of some varieties of Persea americana Peels consumed in Cameroon. Scientific World Journal, vol. 2021, p. 8882594.

Godoy-Gallardo, M., Eckhard, U., Delgado, L. M., de Roo Puente, Y.J.D., Hoyos-Nogués, M., Gil, F.J., & Perez, R.A., 2021. Antibacterial approaches in tissue engineering using metal ions and nanoparticles: From mechanisms to applications. Bioactive Materials, vol. 6, no. 12, pp. 4470–4490.

González, P., Lozano, P., Ros, G., Solano, F., 2023. Hyperglycemia and oxidative Stress: An integral, updated and critical overview of their metabolic interconnections. International Journal of Molecular Sciences, vol. 24, p. 9352.

Griffin, S.P., & Bhagooli, R., 2004. Measuring antioxidant potential in corals using the FRAP assay. Journal of Experimental Marine Biology and Ecology, vol. 302, no. 2, pp. 201–211.

Hochma, E., Yarmolinsky, L., Khalfin, B., Nisnevitch, M., Ben-Shabat, S., Nakonechny, F., 2021. Antimicrobial effect of phytochemicals from edible plants. Processes, vol. 9, p. 2089.

Hossain, F., Mostofa, M.G., Alam, A.K., 2021. Traditional uses and pharmacological activities of the genus leea and its phytochemicals: A review. Heliyon, vol. 7, no. 2, p. e06222.

Ilyasov, I.R., Beloborodov, V.L., Selivanova, I.A., Terekhov, R.P., 2020. ABTS/PP decolorization assay of antioxidant capacity reaction pathways. International Journal of Molecular Sciences, vol. 21, no. 3, p. 1131.

Jomova, K., Raptova, R., Alomar, S.Y., Alwasel, S.H., Nepovimova, E., Kuca, K., Valko, M., 2023. Reactive oxygen species, toxicity, oxidative stress, and antioxidants: chronic diseases and aging. Archives of Toxicology, vol. 97, pp. 2499–2574.

Joshi, A., Prasad, S.K., Joshi, V.K., Hemalatha, S., 2016. Phytochemical standardization, antioxidant, and antibacterial evaluations of Leea macrophylla: A wild edible plant. Journal of Food and Drug Analysis, vol. 24, no. 2, pp. 324–331.

Lam, T.P., Tran, N.N., Pham, L.D., Lai, N.V., Dang, B.N., Truong, N.N., Nguyen-Vo, S.K., Hoang, T.L., Mai, T.T., Tran, T.D., 2024. Flavonoids as dual-target inhibitors against α-glucosidase and α-amylase: A systematic review of in vitro studies. Natural Products and Bioprospecting, vol. 14, no. 1, p. 4.

Larit, F., León, F., Benyahia, S., Cutler, S.J., 2019. Total phenolic and flavonoid content and biological activities of extracts and isolated compounds of Cytisus villosus Pourr. Biomolecules, vol. 9, no. 11, p. 732.

Martinez-Gonzalez, A.I., Díaz-Sánchez, Á.G., de la Rosa, L.A., Bustos-Jaimes, I., Alvarez-Parrilla, E., 2019. Inhibition of α-amylase by flavonoids: Structure activity relationship (SAR). Spectrochimica Acta Part A: Molecular and Biomolecular Spectroscopy, vol. 206, pp. 437–447.

Mohamed, E.A.H., Siddiqui, M.J.A., Ang, L.F., Sadikun, A., Chan, S.H., Tan, S.C., Yam, M.F., 2012. Potent α-glucosidase and α-amylase inhibitory activities of standardized 50% ethanolic extracts and sinensetin from Orthosiphon stamineus Benth as anti-diabetic mechanism. BMC Complementary and Alternative Medicine, vol. 12, no. 1, pp. 176–189.

Muflihah, Y.M., Gollavelli, G., Ling, Y.C., 2021. Correlation study of antioxidant activity with phenolic and flavonoid compounds in 12 indonesian indigenous herbs. Antioxidants (Basel), vol. 10, no. 10, p. 1530.

Murugaiyan, J., Kumar, P.A., Rao, G.S., Iskandar, K., Hawser, S., Hays, J.P., Mohsen, Y., Adukkadukkam, S., Awuah, W.A., Jose, R.A.M., Sylvia, N., Nansubuga, E.P., Tilocca, B., Roncada, P., Roson-Calero, N., Moreno-Morales, J., Amin, R., Kumar, B.K., Kumar, A., Toufik, A.R., Zaw, T.N., Akinwotu, O.O., Satyaseela, M.P., van Dongen, M.B.M., 2022. Progress in alternative strategies to combat antimicrobial resistance: focus on antibiotics. Antibiotics (Basel), vol. 11, no. 2, p. 200.

Ngan, L.T., Moon, J.K., Kim, J.H., Shibamoto, T., Ahn, Y.J., 2012. Growth-inhibiting effects of Paeonia lactiflora root steam distillate constituents and structurally related compounds on human intestinal bacteria. World J Microbiol Biotechnol, vol. 28, no. 4, pp. 1575–1583.

Oluwagunwa, O.A., Alashi, A.M., Aluko, R.E., 2021. Inhibition of the in vitro Activities of α-amylase and pancreatic lipase by aqueous extracts of Amaranthus viridis, Solanum macrocarpon and Telfairia occidentalis leaves. Frontiers in Nutrition, vol. 8, p. 772903.

Patra, S., Panda, P.K., Panigrahi, D.P., Praharaj, P.P., Bhol, C.S., Mahapatra, K.K., Bhutia, S.K., 2019. Terminalia bellirica extract induces anticancer activity through modulation of apoptosis and autophagy in oral squamous cell carcinoma. Food and Chemical Toxicology, vol. 136, p. 111073.

Piaru, S.P.; Mahmud, R.; Majid, A.M.S.A.; Nassar, Z.D.M., 2012. Antioxidant and antiangiogenic activities of the essential oils of Myristica fragrans and Morinda citrifolia. Asian Pacific Journal of Tropical Medicine, vol. 5, pp. 294–298.

Saleem, M., Durani, A.I., Asari, A., Ahmed, M., Ahmad, M., Yousaf, N., Muddassar, M., 2023. Investigation of antioxidant and antibacterial effects of citrus fruits peels extracts using different extracting agents: Phytochemical analysis with in silico studies. Heliyon, vol. 9, no. 4, p. e15433.

Singh, S., Singh, R.P., 2008. In vitro methods of assay of antioxidants: an overview. Food Reviews International, vol. 24, no. 4, pp. 392–415.

Uddin, S., Bin Safdar, L., Fatima, I., Iqbal, J., Ahmad, S., Ahsan Abbasi, B., & Masood Quraishi, U., 2022. Bioprospecting roots, stem and leaves extracts of Berberis baluchistanica Ahrendt. (Berberidaceae) as a natural source of biopharmaceuticals. Journal of Taibah University for Science, vol. 16, no. 1, pp. 954–965.

Ullah, A., Munir, S., Badshah, S.L., Khan, N., Ghani, L., Poulson, B.G., Emwas, A.H., Jaremko, M., 2020. Important flavonoids and their role as a therapeutic agent. Molecules, vol. 25, no. 22, p. 5243.

Umamaheswari, M., Chatterjee T.K., 2007. In vitro antioxidant activities of the fractions of Coccinnia grandis L. leaf extract. African Journal of Traditional, Complementary and Alternative Medicines, vol. 5, no. 1, pp. 61–73.

Vo, V.C., 2021. Dictionary of Vietnamese medicinal plants, volume 1. Hanoi Medical Publishing House, Vietnam, 1677 pages.

Xie, J., Schaich, K.M., 2014. Re-evaluation of the 2,2-diphenyl-1-picrylhydrazyl free radical (DPPH) assay for antioxidant activity. Journal of Agricultural and Food Chemistry, vol. 62, no. 19, pp. 4251–4260.

Zakhele, M.D., Chiy-Rong, C., Bongani, S.D., Chi-I, C., 2023. In vitro antioxidant, antiglycation, α-glucosidase and α-amylase inhibitory activities of extracts and solvent fractions of Elaeocarpus serratus L. Biocatalysis and Agricultural Biotechnology, vol. 52, p. 102827.

Zhang, L.L., Zhang, L.F., Xu, J.G., 2020. Chemical composition, antibacterial activity and action mechanism of different extracts from hawthorn (Crataegus pinnatifida Bge.). Scientific Reports, vol. 10, no. 1, p. 8876.

